# The magnitude and durability of neutralizing antibody responses to human papillomavirus vaccine do not depend on DNA sensing pathways

**DOI:** 10.1101/2025.10.31.685858

**Authors:** Juhi Patel, Zida Yang, Sara Ping, Trevor W. Simon, Jesse J. Waggoner, Erin M. Scherer

## Abstract

How human papillomavirus (HPV) vaccines elicit robust and enduring neutralizing antibody (nAb) responses is unknown, yet such information is valuable to vaccinology. In addition to the major capsid protein, L1, the 4-valent HPV vaccine reportedly also contains recombinant L1 DNA. Nucleic acids in vaccines can enhance antibody responses, but whether L1 DNA enhances HPV vaccine antibody responses is understudied. We tested whether 9-valent HPV (9vHPV) vaccine lots contained L1 DNA and if peak or long-term 9vHPV vaccine-elicited nAb responses were dependent on DNA sensors TLR9, AIM2, or cGAS, or STING or MyD88, the adaptors for cGAS and TLR9, respectively. To quantify L1 DNA, we extracted total nucleic acid per 9vHPV dose and applied singleplex quantitative PCR to amplify HPV6/11/16/18/31/33/45/52/58 L1 sequences. Wildtype mice (wt; C57BL/6J) or mice deficient in DNA sensing pathways (TLR9^-/-^, AIM2^-/-^, cGAS^-/-^, STING^gt/gt^, MyD88^-/-^) were administered 9vHPV vaccine or equivalent adjuvant only at 0, 4, and 12 weeks. We measured serum HPV16 and HPV18 nAb titers at 16, 26, 40, and 53-56 weeks and HPV16- and HPV18-specific bone marrow plasma cell responses at 53-56 weeks. We detected 5-44 and 104-351 copies of HPV6 and HPV18 L1 DNA per 9vHPV vaccine dose, but no other types (n=3 lots). We found no consistent difference in geometric mean nAb titers between mice strains that received 9vHPV at any time point tested or in median HPV16- or HPV18-specific plasma cell frequencies. Thus, we conclude the magnitude and durability of nAb responses to 9vHPV vaccination do not depend on DNA sensing pathways.

## INTRODUCTION

Antibodies play a key role in vaccine-induced prevention of infectious diseases (1, 2). Because live attenuated vaccines may not be feasible for all pathogens or recipients, the development of subunit vaccines that elicit potent and durable antibody responses is essential. However, knowledge of how to reliably design subunit vaccines that elicit long-lived antibody responses is lacking.

HPV vaccines are examples of subunit vaccines that induce robust humoral immunity for many years with extremely slow antibody decay rates (3–8). HPV vaccine-induced antibody durability is superior to that of many other approved inactivated and subunit vaccines (8–17). Thus, HPV vaccines represent a model for studying how lasting subunit vaccine humoral immunity is generated. The 9-valent HPV (9vHPV) vaccine is the only licensed HPV vaccine currently available in the US. It protects against HPV types that commonly cause genital warts (HPV 6 and 11) and cancer (HPV 16, 18, 31, 33, 45, 52, and 58) (18–22). HPV vaccines are comprised of virus-like particles (VLP), where each HPV type VLP assembles from 360 units of its major capsid protein, L1 (23–25). L1 encapsidates recombinant DNA plasmids similar to its eight kilobase double-stranded DNA genome, and the US Food and Drug Administration (US FDA) reported that the predecessor of the 9vHPV vaccine, the 4-valent HPV vaccine, contains recombinant L1 DNA (26–29).

Antigen-associated DNA enhances antibody, germinal center B cell, and plasmablast responses in animal models and has been investigated as an adjuvant (30–33). DNA is sensed by various pattern recognition receptors (PRRs), such as Toll-like receptor 9 (TLR9), which senses hypomethylated DNA CpG sites in the endosomal compartment, and cyclic GMP-AMP synthase (cGAS) and absent in melanoma 2 (AIM2), which sense double-stranded DNA in the cytosol (34, 35). HPV 16 and 18 L1 open reading frames (ORFs) have several CpG sites (36). However, whether HPV vaccine elicited antibody responses depend on DNA sensing pathways has not been studied to our knowledge. Thus, we tested whether we could detect HPV L1 DNA in 9vHPV vaccine lots and if so, whether peak and long-lived 9vHPV-elicited antibody responses depended on DNA sensing pathways.

## RESULTS

### Measuring HPV L1 DNA levels in different 9vHPV vaccine lots

We first sought to confirm whether HPV L1 DNA could be detected in 9vHPV vaccine. To do this, nucleic acids were extracted from single doses (500 µL each) of three separate 9vHPV vaccine lots. Individual 9vHPV doses were divided in half (250 µL x 2) prior to extraction to generate technical replicates, and singleplex quantitative PCR reactions performed on each replicate to amplify HPV 6, 11, 16, 18, 31, 45, 52, or 58 L1 DNA with primers/probes targeting naturally occurring HPV L1 sequences (37). As shown in **Table 1**, we reproducibly detected HPV 6 and 18 L1 DNA in each of three 9vHPV lots at estimated copy numbers of 25-44 or 104-351 per 500 µL 9vHPV dose, respectively. Other types were not detected. This confirmed that HPV L1 DNA was present in 9vHPV vaccine.

**Table 1.** Magnitude of HPV L1 DNA detected in 9vHPV vaccine lots.

|  | **9vHPV**<br>**vaccine lot 1** | **9vHPV**<br>**vaccine lot 2** | **9vHPV**<br>**vaccine lot 3** | **Median** |
| --- | --- | --- | --- | --- |
| **HPV6 L1 DNA** | 25* | 5 | 44 | 25 |
| **HPV18 L1 DNA** | 104 | 351 | 191 | 191 |
| **All other 9vHPV-type L1**<br>**DNA** | ND^ | ND | ND | - |
| *copies/9vHPV vaccine dose<br>^ND, not detected |  |  |  |  |

### Determining a 9vHPV vaccine dose for the mouse model

Prior to testing whether peak and long-lived 9vHPV-elicited antibody responses depended on DNA sensing pathways in the mouse model, we first needed to select a vaccine dose for mice. We tested different dose volumes of 9vHPV vaccine in five groups of male (n=5) and female (n=5) wildtype (C57BL/6J) mice: approximately 1/4 (120 μL), 1/8 (60 μL), 1/16 (30 μL), 1/32 (20 μL), and 1/64 (10 μL) of the human dose volume (**Fig. 1A**). According to the 9vHPV vaccine prescribing information, this corresponds to 4.8-14.4 µg HPV L1 protein for the highest dose or 0.4-1.2 µg HPV L1 protein for the lowest dose, exact amount dependent on HPV type. These 9vHPV dose volumes were administered intramuscularly (i.m.) at Day 0, Week 4, and Week 11 in each study group (**Fig. 1A**) to mimic the adult human 9vHPV dose schedule that is given at 0, 2, and 6 months. As shown in **Figure 1B**, when HPV16 neutralizing antibody (nAb) titers were assessed in sera collected approximately one month after the final dose at Week 16 by pseudovirus neutralization assay, there was no difference in titers by dose volume. However, both the 10 µL and 30 µL dose volumes generated significantly lower HPV18 nAb titers compared to the highest dose volume. Thus, we selected the 60 µL 9vHPV dose volume as the lowest 9vHPV dose volume that did not elicit inferior nAb responses at Week 16. As shown in **Figure 1C**, HPV16 nAb titers peaked approximately one month after the second 60 µL 9vHPV dose at Week 8, and stayed elevated approximately one month after the final 60 µL 9vHPV dose at Week 16. Moreover, male and female mice generated similar HPV nAb titers to the 9vHPV vaccine (**Fig. S1**).

**Figure 1.**
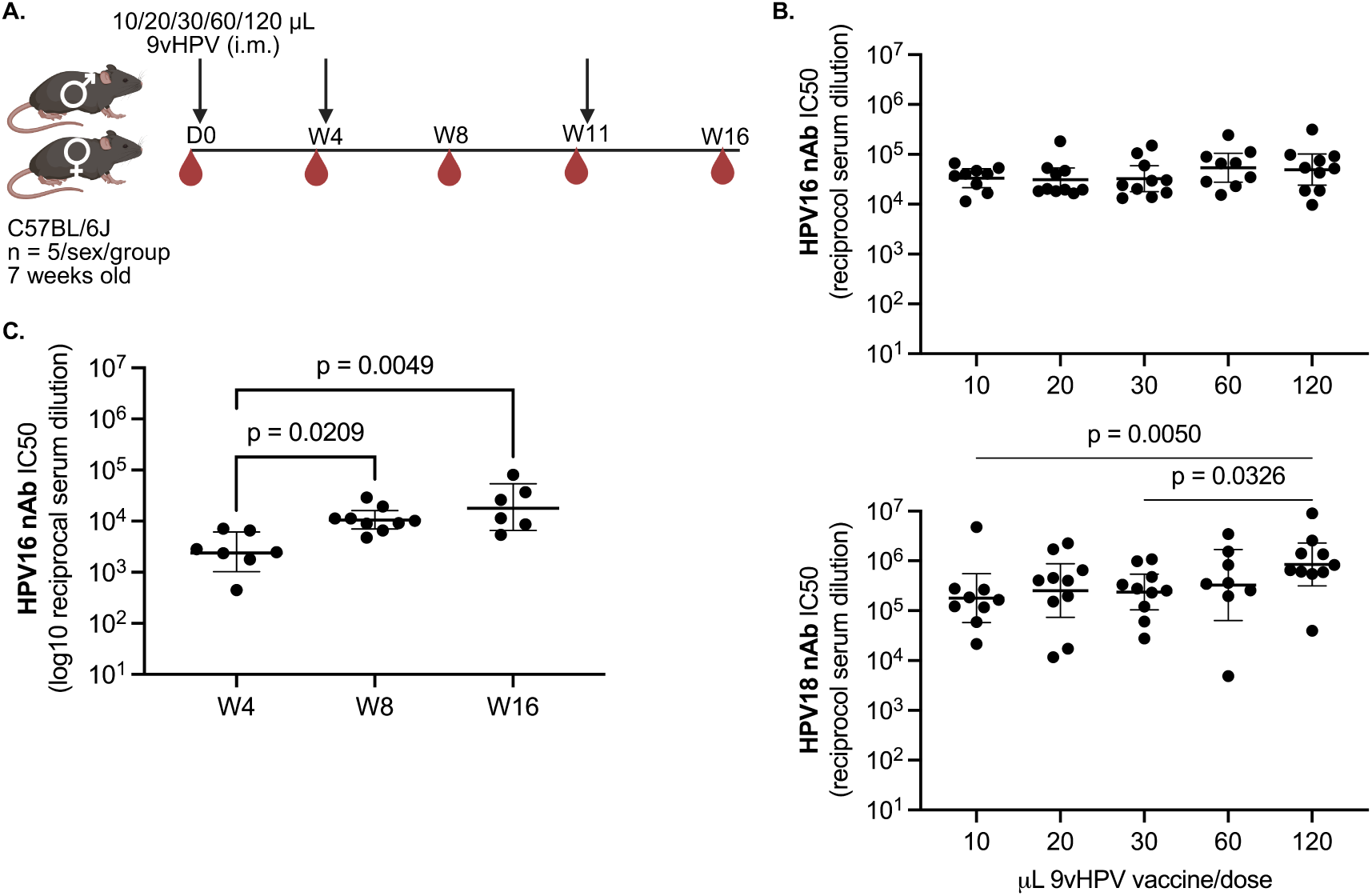
9vHPV vaccine dose titration to identify a dose volume for mice. **A.** Study schematic (D, Study Day; W, Study Week; i.m., intramuscularly) created with BioRender. **B.** Peak serum HPV16 and HPV18 neutralizing antibody (nAb) titers approximately one month after the third and final dose at Week 16 in mice (n = 8-10 per group) that received the indicated dose volumes; 60 *μ*L was selected as the lowest volume that elicited non-inferior HPV16 and HPV18 nAb responses at Week 16. **C.** Serum HPV16 nAb titers approximately one month after the first, second, or third 60 *μ*L 9vHPV vaccine dose (n = 6-9 mice per time point). Geometric mean with 95% confidence intervals shown, where each point represents the result from an individual mouse; Kruskal-Wallis with Dunn’s post-test between dose groups.

### Testing whether peak 9vHPV vaccine-elicited antibody responses depend on DNA sensing pathways

To test whether peak 9vHPV-elicited antibody responses depend on DNA sensing pathways, we administered three, 60 µL 9vHPV doses at Day 0, Week 4, and Week 12 in groups of wildtype mice (male n=5; female n=5), as well as mice globally deficient in TLR9, cGAS, and AIM2 DNA sensing pathways (TLR9^-/-^, cGAS^-/-^ (38), AIM2^-/-^ (39); **Fig. 2A**). We included mice globally deficient in myeloid differentiation factor 88 (MyD88) and stimulator of interferon genes (STING) as controls (MyD88^-/-^ (40) and STING^gt/gt^ (41); **Fig. 2A**), because TLR9 and cGAS signal through these adaptors, respectively. We also included mice globally deficient in TLR4 (TLR4^del/del^ (42, 43); **Fig. 2A)**, as prior work has reported conflicting results on the importance of this pattern recognition receptor pathway to antibody responses elicited by HPV virus-like particles in mice (44, 45). In addition to 9vHPV, we also administered three doses of equivalent adjuvant only as a negative control (amorphous aluminum hydroxyphosphate sulfate, AAHS; 60 µg) at Day 0, Week 4, and Week 12 to separate groups of the same strains of mice (**Fig. 2A**). Serum from mice that received AAHS did not exhibit HPV neutralizing activity one-month post-third/final dose at Week 16 as expected, given that these mice received no HPV L1 antigen (**Fig. 2B**), whereas serum from mice that received 9vHPV exhibited potent HPV16 and HPV18 neutralizing activity (**Fig. 2B**). However, no differences in HPV16 or HPV18 nAb titers were observed between wildtype mice and TLR9^-/-^, cGAS^-/-^, AIM2^-/-^, MyD88^-/-^, STING^gt/gt^, or TLR4^del/del^ mice (**Fig. 2B**), indicating that the magnitude of nAb responses to the 9vHPV vaccine does not depend on DNA sensing pathways or other pathways tested. Furthermore, as before, male and female mice of all strains elicited similar nAb titers to the 9vHPV vaccine (**Fig. S2**).

**Figure 2.**
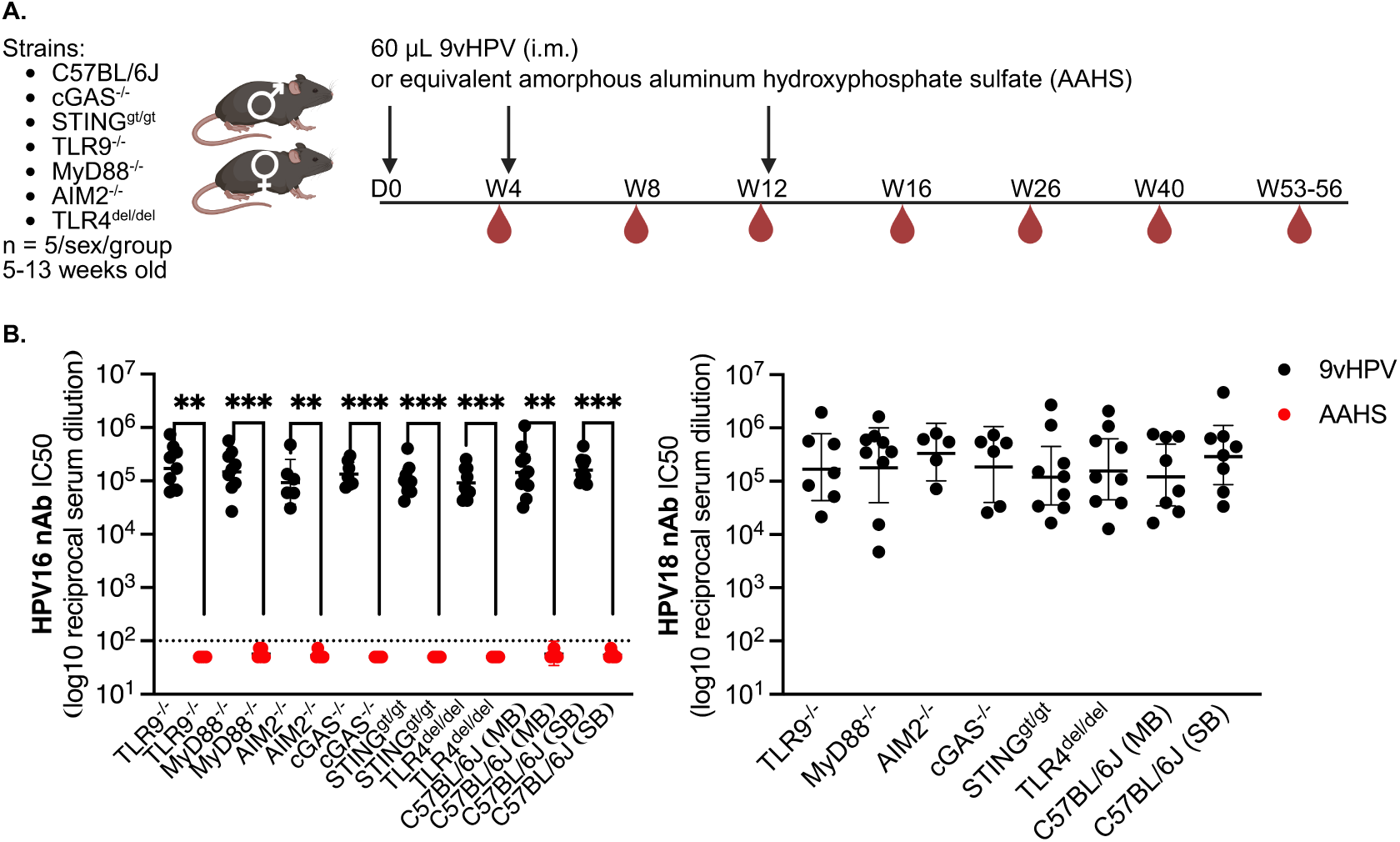
The magnitude of peak nAb responses to the 9vHPV vaccine does not depend on DNA sensing pathways. **A.** Study schematic created with BioRender. **B.** Peak serum HPV16 or HPV18 nAb titers approximately one month after the third and final dose at Week 16 in mice that received 9vHPV (n = 5-9 per group), which contains AAHS as an adjuvant, or equivalent AAHS (60 μg) only (n = 3-8 per group). MB or SB represent maximum barrier or standard barrier vivarium rooms where mice were housed. Geometric mean with 95% confidence intervals shown, where each point represents the result from an individual mouse; Kruskal-Wallis with Dunn’s post-test comparing similarly housed strains to control (wildtype C57BL/6J mice); Mann-Whitney U test between 9vHPV and AAHS groups for a given strain (**p=0.0022-0.007; ***p=0.0002-0.001).

Given that we detected HPV18 L1 DNA in all lots of 9vHPV vaccine tested, including the ones used in mice, but not HPV16 L1 DNA, we also evaluated whether peak HPV18 nAb titers were higher in magnitude than HPV16 nAb titers in any mice strain tested (**Fig. S3**). We found no difference between HPV16 or HPV18 nAb titers at Week 16 for any mice strain that received 9vHPV vaccine. However, n.b., that lack of HPV16 L1 DNA detection in **Table 1** does not necessarily mean that HPV16 L1 DNA is not present in 9vHPV vaccine; it simply means we did not detect it with our methods.

### Testing whether long-term 9vHPV vaccine-elicited antibody responses depend on DNA sensing pathways

To test whether long-term 9vHPV-elicited antibody responses depend on DNA sensing pathways, we assessed HPV neutralizing activity in sera collected at Week 26, 40, and 53-56 after the first 9vHPV vaccine dose (**Fig. 3A-C**). HPV16 neutralizing activity was assessed at all time points, but due to limited amounts of serum collected by submandibular (facial vein) bleeding at Week 26 and 40, HPV18 neutralizing activity was only assessed at Week 53-56 (i.e., **Fig. 3C**), when more blood was collected by cardiac puncture. We found long-term nAb responses to the 9vHPV vaccine did not depend on DNA sensing pathways, MyD88, or TLR4 (**Fig. 3A-D**). However, it should be noted that global functional knockout of STING significantly improved HPV18 nAb titers by 4.1-fold at Week 53-56 (**Fig. 3C**). In addition, we found no consistent differences in HPV nAb titers between sexes within a strain or between the same sex of different strains at any of the time points tested (**Fig. S4**). Moreover, when decay rates of log10 HPV16 nAb titers over time were fitted with a linear regression model for each mouse with results from Week 16, 26, 40, and 53-56, we observed no difference in nAb titer decay rates across strains (**Fig. S5**). These data thus indicate that the durability of nAb responses to the 9vHPV vaccine does not depend on DNA sensing pathways or other pathways tested.

**Figure 3.**
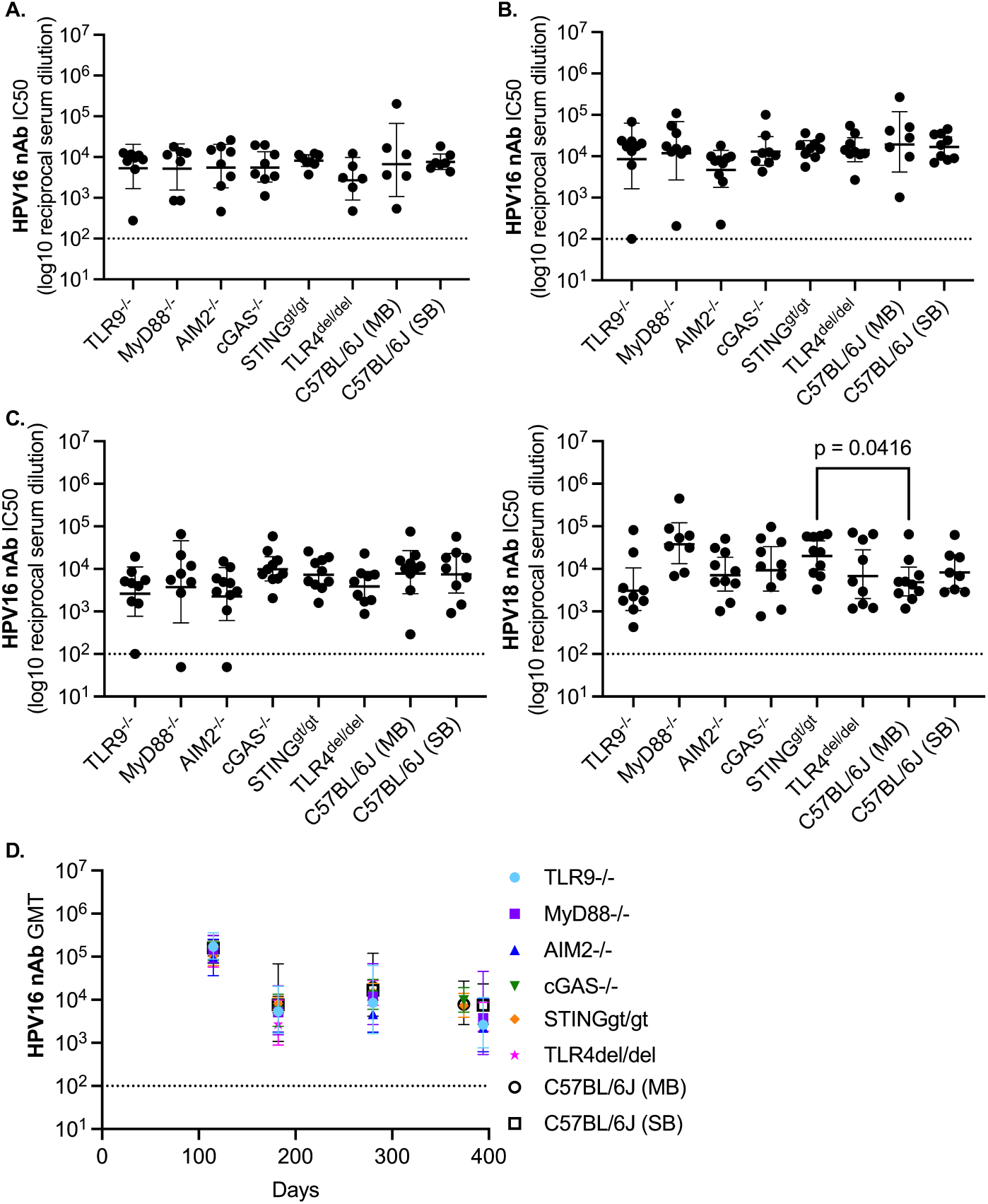
The durability of nAb responses to the 9vHPV vaccine also does not depend on DNA sensing pathways. **A-B.** Serum HPV16 nAb titers in mice that received 9vHPV, approximately six months after the first 9vHPV vaccine dose at Week 26 (**A**; n = 6-8 per group), or approximately nine months after the first 9vHPV vaccine dose at Week 40 (**B**; n = 7-10 per group). **C.** Serum HPV16 or HPV18 nAb titers approximately one year after the first 9vHPV vaccine dose at Week 53-56 (n = 8-10 per group). Geometric mean with 95% confidence intervals shown, where each point represents the result from an individual mouse; Kruskal-Wallis with Dunn’s post-test comparing similarly housed strains to control. **D.** Change in geometric mean titers (GMT) with 95% confidence intervals of HPV16 nAb responses over time.

### Assessing whether long-lived, 9vHPV vaccine-elicited bone marrow plasma cell responses depend on DNA sensing pathways

As steady state antibody levels in blood are replenished by long-lived plasma cells in the bone marrow (46–48), we sought to determine if there were differences in the frequencies of HPV-specific plasma cells in bone marrow approximately a year after the first 9vHPV vaccine dose between wildtype mice and TLR9^-/-^, cGAS^-/-^, AIM2^-/-^, MyD88^-/-^, STING^gt/gt^, or TLR4^del/del^ mice. To do this, we developed an ELISPOT that measured the frequency of HPV16- or HPV18-specific IgG secreting cells among total IgG-secreting cells in bone marrow (**Fig. 4A**). To detect HPV16- or HPV18-specific IgG secreting cells, we coated ELISPOT plates with purified HPV16 or HPV18 virus-like particles containing L1 and L2, the minor capsid protein (**Fig. 4A**). We found a significantly lower frequency of HPV18-specific IgG plasma cells in mice that received AAHS versus those receiving 9vHPV vaccine, as expected (**Fig. 4B**). However, we observed no difference in the frequency of HPV16- or HPV18-specific IgG plasma cells between mice strains receiving 9vHPV vaccine approximately a year after the first dose (**Fig. 4B**), further supporting the nAb results that 9vHPV vaccine-elicited B cell responses do not depend on DNA sensing pathways.

**Figure 4.**
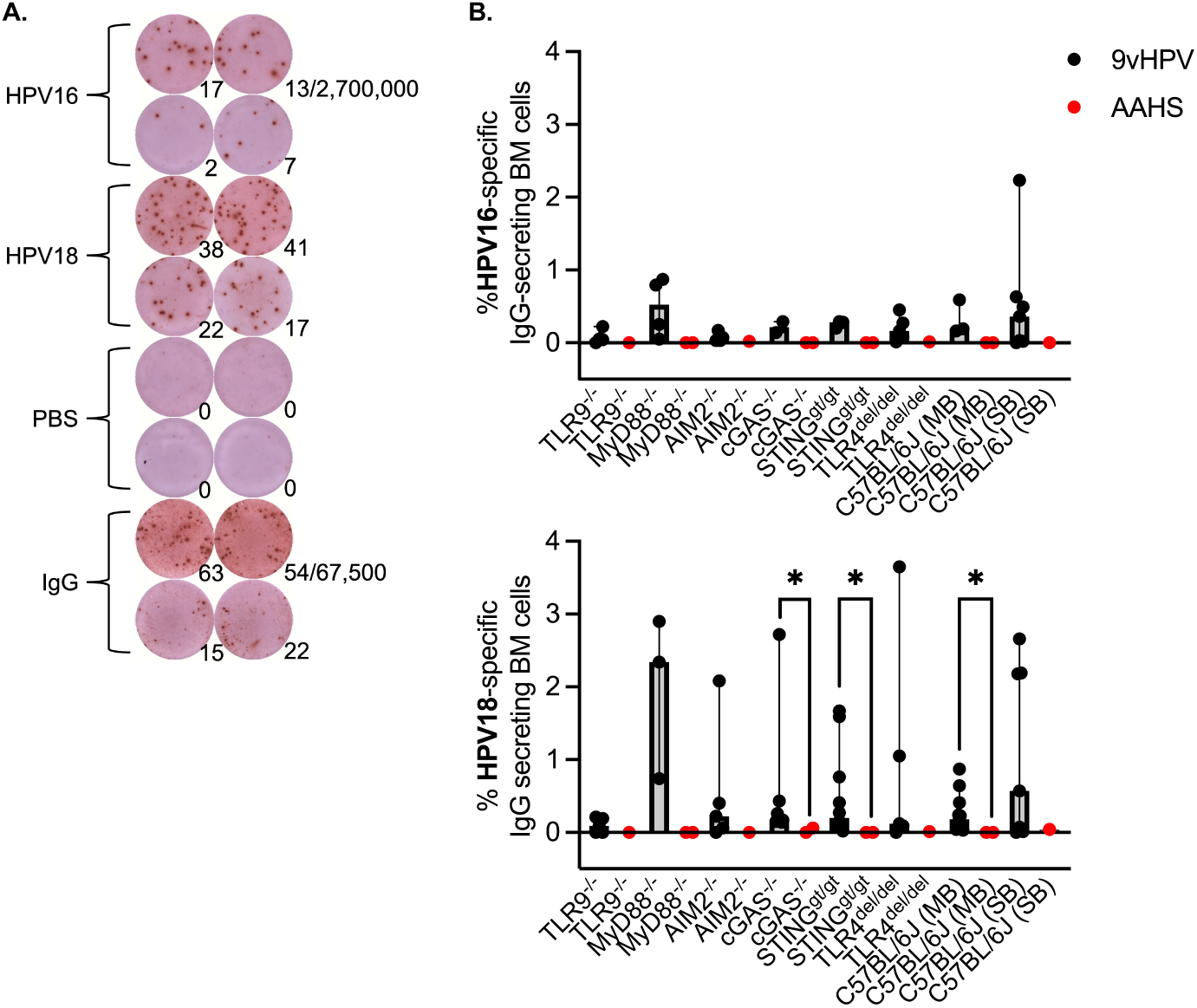
HPV-specific plasma cell responses to the 9vHPV vaccine do not depend of DNA sensing pathways. **A.** Representative image of ELISPOT plate wells coated with HPV16 or HPV18 virus-like particles (VLPs), VLP vehicle (phosphate buffered saline, PBS), or anti-mouse IgG. Numbers at bottom right of each well indicate spot forming units (SFUs) per well. Denominators indicate number of input cells plated in top dilution row in duplicate (2,700,000 bone marrow (BM) cells for VLP or PBS wells; 67,500 BM cells for IgG wells), where bottom row is 3-fold dilution of cells. **B.** Frequencies of HPV16- or HPV18-specific BM cells of total IgG-secreting BM cells as determined by ELISPOT approximately one year after the first 9vHPV vaccine dose at Weeks 53-56 (n = 2-7 per group HPV16/9vHPV; n = 1-2 per group HPV16/AAHS; n = 3-10 per group HPV18/9vHPV; n = 1-2 per group HPV18/AAHS). Median with 95% confidence intervals shown, where each point represents the result from an individual mouse; Kruskal-Wallis with Dunn’s post-test comparing similarly housed strains to control. Mann-Whitney U test between 9vHPV and AAHS groups for a given strain (*p=0.0303-0.0444).

### Estimating the relationship between HPV nAb responses in blood and HPV-specific plasma cell responses in bone marrow

To further examine the relationship between nAb titers and bone marrow plasma cell frequencies, we estimated the correlation between HPV16 or HPV18 nAb titers in sera and the frequency of HPV16 or HPV18-specific IgG plasma cells in bone marrow across all mice with paired specimens that received 9vHPV (**Fig. 5**). We found strong and highly significant positive relationships between these immune parameters for both HPV16 and HPV18 (**Fig. 5**), as expected given the crucial antibody secreting role of plasma cells in the bone marrow.

**Figure 5.**
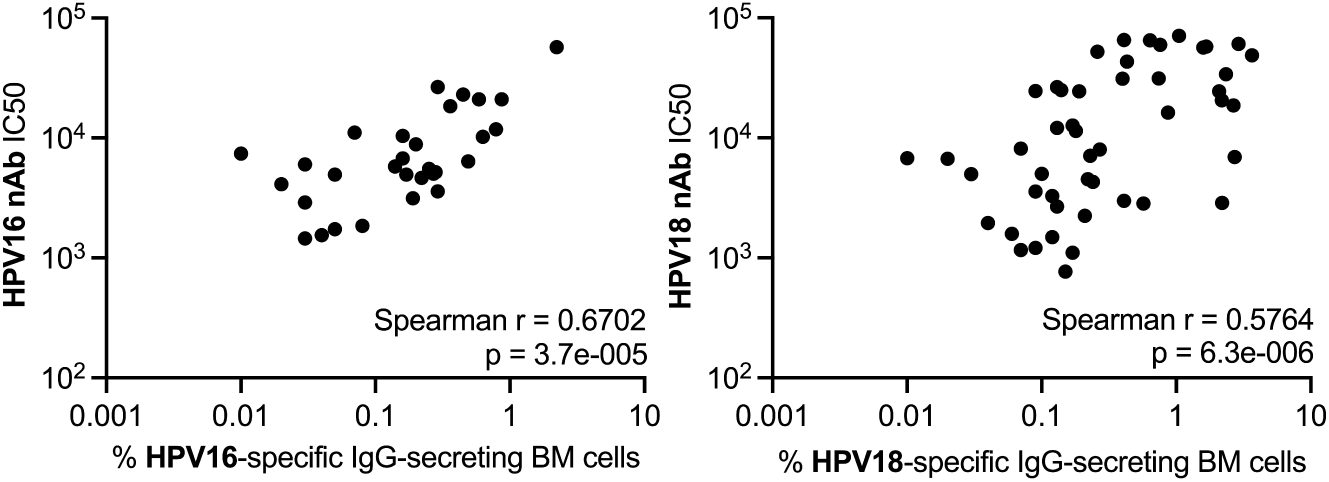
HPV nAb titers correlate with HPV-specific IgG-secreting bone marrow (BM) plasma cell frequencies one year post-first 9vHPV dose. Scatter plots of HPV16 or HPV18 nAb titers and frequencies of HPV16- or HPV18-specific IgG-secreting BM cells, respectively, approximately one year after the first 9vHPV vaccine dose at Weeks 53-56. Each point represents the result from an individual mouse (n = 31 for HPV16; n = 53 for HPV18); Correlation estimated between immune parameters with Spearman coefficients.

## DISCUSSION

In 2011, the US FDA acknowledged that the 4vHPV vaccine contains HPV L1 DNA as an expected result of the manufacturing process that poses no safety concerns (26). As for many protein-based biologics (e.g., subunit vaccines, monoclonal antibodies, etc.), recombinant DNA is required to express the L1 proteins that are purified for incorporation into the vaccine. Here, we asked whether HPV L1 DNA was also present in the updated 9vHPV vaccine and if so, whether DNA sensing pathways were contributing to the remarkable magnitude and durability of HPV vaccine-elicited antibody responses. We confirmed the presence of HPV L1 DNA in multiple vaccine lots, specifically HPV6 and HPV18 L1 DNA. However, to our surprise, both the magnitude and durability of 9vHPV-elicited nAb responses in mice do not appear to depend on DNA sensing pathways tested, specifically TLR9, cGAS, or AIM2. We also show here that the magnitude and durability of 9vHPV-elicited nAb responses do not appear to depend on TLR4 or other TLR pathways that signal through MyD88, including potentially TLR2 that can detect yeast cell wall components (49–51). This is relevant because both the 4vHPV and 9vHPV vaccines are produced in baker’s yeast (*Saccharomyces cerevisiae*) (52, 53).

In contrast to our results, a prior study by Yang *et al.* found that antibody responses to HPV16 VLPs were lower in MyD88^-/-^ mice and TLR4-hyporesponsive mice compared to wildtype mice (45). Yet, a subsequent study by Thönes *et al.* did not reproduce the TLR4 result (44). Specifically, Yang *et al.* found that global knockout of MyD88 eliminated IgG(1–3) responses to HPV16 VLPs (45), in contrast to our finding that global MyD88 knockout had no impact in nAb titers to 9vHPV. There are many differences between this prior study and ours that could account for these disparate results: first, we analyzed serum responses via the gold standard HPV pseudovirus neutralization assay that correlates strongly with binding IgG responses in both mice (44) and humans (54), whereas Yang *et al.* measured serum IgG subclass binding to HPV16 VLPs via ELISA (45); second, we immunized mice via the same route as humans (intramuscularly) whereas Yang *et al.* immunized mice intravenously (45); third, our antigen (60 µL 9vHPV) is equivalent to ∼7.2 µg HPV16 L1 VLPs expressed from yeast with AAHS adjuvant, whereas the antigen in Yang *et al.* was 10 µg HPV16 L1 VLPs expressed from baculovirus/insect cells with no adjuvant (45); fourth, we immunized mice on Day 0, 28, and 84, and collected blood on Day 114 onward, whereas Yang *et al.* immunized mice on Day 0, 7, and 14 and collected blood on Day 24 (45); and fifth, the MyD88^-/-^ mice used by Yang *et al.* were maintained by Prof. Shizuo Akira’s group (45), whereas our mice were from Akira’s group further backcrossed to C57BL/6J (40) and purchased from The Jackson Laboratory. Yang *et al.* also observed that global knockout of TLR4 reduced serum IgG3 responses to HPV16 VLPs compared to wildtype mice, as well as serum IgG2a and IgG2b responses (less than IgG3 decrease; no statistical significance shown), but found no change in the predominant IgG1 responses to HPV16 VLPs relative to wildtype mice (45). In this case, the TLR4 mutant mouse strain used also differed from ours (C3H/HeJ mice in the case of Yang *et al.* (45), and TLR4^del/del^ mice (42, 43) here). When Thönes *et al.* conducted a similar study in C3H/HeJ (TLR4 mutant) and C3H/HeOuJ (wildtype) mice – the same mice used by Yang *et al.* – they found no difference in the IgG response to HPV16 VLPs in C3H/HeJ versus C3H/HeOuJ mice (44). In the Thönes *et al.* study, C3H/HeJ or C3H/HeOuJ mice were immunized subcutaneously with 2.5 or 10 µg HPV16 VLPs produced from baculovirus/insect cells with no adjuvant at Day 0 and 14 and with blood collected on Day 24 (44). Serum IgG binding to HPV16 VLPs was assessed by capture ELISA (44); so overall more similar to the Yang *et al.* study. It is not immediately clear what could account for the differences in outcomes in the case of the MyD88^-/-^ mice, but it seems that in the case of the C3H/HeJ mice, the reduction in IgG3, 2a, and 2b responses to VLPs was insufficient to diminish the total IgG response (44), which appeared to be predominantly IgG1 (45). It should also be noted that while the nAb responses to 9vHPV vaccine appear to be durable from approximately month 6 to month 13 in mice – which was not known prior to this study – we do observe a decay from the peak at month 4 to month 6, which along with the antibody response kinetics following the first, second, and third doses, mimics HPV vaccine-elicited antibody kinetics observed in humans (4, 6, 7, 55, 56). However, murine studies of antibody responses to viral infections reveal antibody responses that do not decay from the peak (48), highlighting crucial differences in the immune responses elicited to viral infections and subunit vaccines in mice that we are keen to explore further.

Limitations of our study include the following: First, a priori we did not know the L1 DNA sequences utilized to manufacture the vaccines, as they are not publicly available. Therefore, it is possible our RT-PCR primers and probes, which are specific for naturally occurring L1 sequences, are not specific for L1 sequences in the vaccine and thus could not amplify them (e.g., if L1 sequence modifications were made to improve protein expression). Second, given the number of mice followed long-term, it was not possible to replicate this pilot study. Third, we observed batch effects in neutralization assays, which can be avoided in future studies by testing all specimens from the same mouse together. This was not feasible for this study, and our priority was to determine if there were any difference in nAb titers between strains at any given time point; thus all specimens from a given time point were tested simultaneously. Fourth, we did not also attempt to remove residual DNA in 9vHPV vaccine (e.g., by applying DNase to 9vHPV doses), due to the risk of impacting the structural integrity of the VLPs, which bind to DNA. Fifth, we did not test mice globally deficient in mitochondrial antiviral signaling (MAVS) (57), which is a global adaptor for double-stranded RNA retinoic acid-inducible gene I (RIG-I)-like sensors RIG-I and melanoma differentiation-associated protein-5 (MDA5) (58). However, double-stranded DNA viruses such as herpes, poxviruses, and adenoviruses, can activate RLRs (59). Therefore, this remains a potential future direction.

Other potential future directions include repeating this work in a smaller number of mice strains (e.g., AIM2^-/-^, MyD88^-/-^, STING^gt/gt^, and wt mice), without nAb testing batch effects, in mice receiving only one or two doses of 9vHPV vaccine. The rationale being that fewer than three HPV vaccine doses is currently recommended by the World Health Organization (60), and differences could become apparent with less saturation of the humoral response. We also aim to examine whether peak memory B cell responses to 9vHPV vaccination differ between mice deficient in DNA sensing pathways relative to wt mice in future work, given that memory B cells and antibody-secreting cells have opposing transcriptional programs (61, 62). This was not feasible here, as we followed a large number of mice for over a year. Depending on the results from these future studies, next steps could include 9vHPV studies in conditional knockout mice (e.g., mice where only B cells or dendritic cells are deficient DNA sensing pathways).

## ACKNOWLEDGMENTS

The Scherer Lab would like to thank Karen Hannah and Madeleine Rasche for assistance with experimental work, Rustom Antia for generous and detailed advice on analyzing antibody decay half-lives, Emory Division of Animal Resources for animal care, and Merck for gift of AAHS to Emory University. The authors would like to acknowledge the following BioRender citations: **Fig. 1**: Created in BioRender. Scherer, E. (2025) https://BioRender.com/jvvdz6x **Fig. 2**: Created in BioRender. Scherer, E. (2025) https://BioRender.com/264buux **Fig. 3**: Created in BioRender. Scherer, E. (2025) https://BioRender.com/vfil6mb **Fig. 4**: Created in BioRender. Scherer, E. (2025) https://BioRender.com/zogrjm1 **Fig. 5**: Created in BioRender. Scherer, E. (2025) https://BioRender.com/kyrhdlj **Fig. S4:** Created in BioRender. Scherer, E. (2025) https://BioRender.com/xjy7r7b

## AUTHOR CONTRIBUTIONS

**JP:** Investigation, Writing – Original Draft; **ZY:** Investigation; **SP:** Investigation, Writing – Original Draft, Writing – Reviewing and Editing; **TWS:** Investigation, Writing – Reviewing and Editing; **JJW:** Methodology, Formal Analysis, Writing – Reviewing and Editing, Supervision; **EMS:** Conceptualization, Methodology, Formal Analysis, Resources, Writing – Original Draft, Writing – Reviewing and Editing, Visualization, Supervision, Project Administration, Funding Acquisition.

## MATERIALS AND METHODS

### Quantitative PCR (qPCR)

Published primer and probe sets (37) were used in singleplex reactions for amplification of L1 gene sequences from HPV6, 11, 16, 18, 31, 45, 52, and 58. A new primer-probe set was designed using Primer3 software for HPV33 based on all L1 sequences available in GenBank. Primers were purchased from Integrated DNA Technologies and probes from Biosearch Technologies. All probes but HPV31 were labeled with Cal Fluor Orange 560; HPV31 was labeled with CIV 550. HPV33 primer-probe sequences were evaluated in silica using BLAST (National Center for Biotechnology Information) to check exclusivity. These were then evaluated using plasmids containing HPV L1 sequences to identify the set with the most sensitive detection (earliest Ct values). All primers were used at a concentration of 400 nM in the final reaction, and probes were used at 200 nM. Plasmids were diluted to 10^8^, 10^6^, 10^4^, 10^2^, 10^0^ in molecular grade water. qPCR was performed using a 20 µL reaction with the Luna Universal Probe qPCR Master Mix (New England Biolabs), using 5 µL of eluted DNA and the following cycling conditions: 95°C for 2 minutes, then 45 cycles of 95°C for 15 seconds and 60°C for 60 seconds. qPCR was run with plasmid dilution series to evaluate the limit of detection. All the primer probe sets had strong curves (data not shown) and detected down to 1 copy/µL of plasmids. 10^2^ concentration was selected as a control for the qPCR runs testing eluted 9vHPV vaccine DNA. 9vHPV vaccine doses of 500 µL were split into two 250 µl aliquots and extracted in duplicate using an Apex Kingfisher instrument and the MagMAX viral/pathogen kit. Nucleic acids were eluted in 100 µL of 10 mM Tris-HCl buffer. Five microliters of eluate were then tested in singleplex qPCR for each type. 9vHPV vaccine from three different lots were tested.

### Animal Experiments

All experimental animal procedures were approved by the Institutional Animal Care and Use Committee of Emory University.

#### 9vHPV dose titration

Five male and five female C57BL/6J mice (The Jackson Laboratory (JAX) strain # 000664) at 7 weeks of age were assigned to the following dose groups: 10 µL, 20 µL, 30 µL, 60 µL, or 120 µL 9vHPV vaccine (Lot 1 in **Table 1**). 9vHPV doses were administered i.m. at one to three sites (up to 50 µL per site) in the rear thigh muscle at weeks 0, 4, and 11. Blood samples were collected under isoflurane-induced anesthesia by submandibular (facial) vein at weeks 0, 2, 4, 6, 8, 11, and 14 into serum gel separator tubes (Sarstedt 20.1291). Mice were euthanized by American Veterinary Medical Association (AVMA) approved methods at week 16. Specifically, blood was collected under isoflurane-induced anesthesia by cardiac puncture into serum gel separator tubes (BD Microtainer 365967). and bone marrow was harvested as described below. Sixty microliters of 9vHPV was selected as the optimal dose based on HPV16 and HPV18 neutralization data when analyzed in aggregate for both sexes (i.e., n = 10 mice/dose group). All mice were housed in a maximum barrier facility.

#### Assessing the dependency of DNA-sensing pathways on 9vHPV antibody responses

The following mouse strains were used for this experiment: C57BL/6J; AIM2^-/-^ (B6.129P2-Aim2^Gt(CSG445)Byg^/J; JAX strain # 013144 (39)); TLR9^-/-^ (generated through a custom breeding project at JAX with C57BL/6-Tlr9^em1.1Ldm^/J strain # 034449); STING^gt/gt^ (C57BL/6J-Sting1^gt^/J; JAX strain # 017537 (41)); MyD88^-/-^ (B6.129P2(SJL)-Myd88^tm1.1Defr^/J; JAX strain # 009088 (40)); TLR4^del/del^ (B6.B10ScN-Tlr4^lps-del^/JthJ; JAX strain # 007227 (42, 43)); and cGAS^-/-^ (B6(C)-Cgas^tm1d(EUCOMM)Hmgu^/J; JAX strain # 026554 (38)). C57BL/6J, cGAS^-/-^, and STING^gt/gt^ mice were housed in a maximum barrier facility at Emory, whereas C57BL/6J, AIM2^-/-^, TLR9^-/-^, MyD88^-/-^ and TLR4^del/del^ mice were housed in a standard barrier facility. Five male and five female mice of each strain at 5-13 weeks of age were assigned per group, except C57BL/6J mice, which served as controls in each facility. Mice received three doses of 9vHPV vaccine (60 µL per dose; Lot 3 in **Table 1**) or three doses of amorphous aluminum hydroxyphosphate sulfate (AAHS; 70 µL per dose to yield equivalent AAHS as 60 µL 9vHPV). 9vHPV or AAHS was administered i.m. at 0, 4, and 12 weeks. Blood was collected under isoflurane-induced anesthesia by facial vein at 4, 8, 12, 16, 26, 40, and 53-56 weeks. Mice were euthanized by AVMA approved methods at 53 weeks (C57BL/6J, cGAS^-/-^, and STING^gt/gt^ mice) or 56 weeks (C57BL/6J, AIM2^-/-^, TLR9^-/-^, MyD88^-/-^ and TLR4^del/del^ mice). On those days, blood was collected under isoflurane-induced anesthesia by cardiac puncture and bone marrow was harvested.

### Isolation of bone marrow cells

An incision around the upper thigh was made and skin removed. Muscle from the legs was removed with scissors, ensuring not to break the bones in the legs. The hip joint was then cut from the ilium. Legs were placed in ice-cold R10 (Roswell Park Memorial Institute medium (RPMI; Cytiva SH30255.01 or equivalent), 10% heat inactivated fetal bovine serum (FBS; R&D Systems #S12450H), 1X penicillin-streptomycin (Cellgro or equivalent), 1X L-glutamine (Gibco or equivalent), 0.1% betamercaptoethanol (Sigma Aldrich)) with 2 mM EDTA (Quality Biological).

After collection, legs were rinsed in 70% ethanol (Decon Labs) followed by three consecutive rinses in cold sterile 1X phosphate buffered saline (PBS) (Corning 21-040-CM or equivalent) to remove ethanol from the surface of the legs. Inside a sterile petri dish (VWR), remaining tissue was removed using sterile scalpel and tweezers and the epiphyses of the bones were removed with scalpel. A 10 mL sterile syringe (Air-Tite) equipped with 25-gauge needle and filled R10 with 2 mM EDTA was applied to flush the bone marrow from both ends of the bone shaft into a sterile 70 µm cell filter (Falcon) and 50 mL conical tube (Falcon). The bones were flushed until blanched. The plunger bulb of a sterile 3 mL syringe (Air-Tite) was used to gently press the bone marrow through the filter and the filter was rinsed with R10 with EDTA. After pelleting cells, red blood cells were lysed with ammonium-chloride-potassium (ACK) lysis buffer (Stemcell Technologies) and quenched with R10 with EDTA. Cells were then pelleted and resuspended in R10 with EDTA prior to counting with a Guava Muse cell analyzer in ViaCount buffer (Cytek).

### Isolation of mouse splenocytes

Mice spleens were homogenized by macerating the spleen on a 100 µm mesh filter (Falcon) with the plunger bulb of a 3 mL syringe. Additional R10 media (without EDTA) was used to flush the spleen through the filter into a conical tube. Cells were pelleted and red blood cells lysed with ACK buffer and quenched with R10. The cells were pelleted again, resuspended in R10, filtered through a 70 µm filter into a clean conical tube, and counted with the Guava Muse in ViaCount buffer.

### HPV pseudovirus (psV) and HPV virus-like particles (VLPs)

HPV psV were generated using the same procedures as previously described in Scherer *et al.* (63), except that psV were purified by density ultracentrifugation on 0.7 mL gradients at 41,000 rpm for ∼6 hours at 16°C in a Sw Ti 41 rotor in 13.2 mL sterile ultracentrifuge tubes (Beckman Coulter C14293). HPV VLPs were generated for bone marrow ELISPOTs in the same manner, except no pYSEAP was used during transfection, and 0.25% benzonase (SigmaAldrich EB263) and 0.25% plasmid safe DNAse (Biosearch Technologies E3101K) were used instead of RNAse during maturation. Density gradient fractions containing psV and VLPs were screened by ELISA and psV titered or VLPs quantified by L1 SDS-PAGE densitometry with reference to BSA standards prior to use as described^4^.

### HPV pseudovirus neutralization assay

The same procedure was used as previously described in Scherer *et al.* (63) except that cells were seeded into an Edgewell 96 well plate (Thermo Fisher Scientific 167542). The outside troughs of the plate were filled with 3 mL of 1XPBS before incubating at 37°C for 4 hours. Sera was initially screened at 1/100 starting dilution in duplicate in a threefold dilution series. Potently neutralizing sera were diluted further (e.g., 1/2700 starting dilution) in subsequent experiments prior to analyzing in duplicate.

### Bone marrow ELISPOT

Millipore Multiscreen-IP 96-well plates (MSIPS4510) were coated with antigens 24 hours prior to use. ELISpot plates were washed according to manufacturer recommendations with 35% ethanol, deionized water, and 1X PBS. Plates were coated with donkey anti-mouse IgG antibody (Jackson ImmunoResearch 715-005-150) at 10 µg/mL in 1X PBS, 1X PBS, or HPV16 or HPV18 VLPs at 10 µg/mL 1X PBS. Depending on cell yield, each sample was tested in duplicate for each antigen. Plates were stored at 4°C until use.

After 24 hours, coated plates were washed four times with PBS-T (0.05% (v/v) Tween-20 (Fisher Scientific or equivalent) in 1X PBS) and blocked with R10 with 2 mM EDTA at 37°C for 1-2 hours. After blocking, R10 with EDTA was added to all wells. Two to four million bone marrow cells were then added to the first well of HPV VLPs or PBS wells; and 50,000-100,000 bone marrow cells were added to the first well of the IgG wells. Then one three-fold serial cellular dilution was performed and plates incubated at 37°C for approximately 18 hours.

After ∼18 hours, plates were washed with 1X PBS and then with PBS-T. Biotinylated anti-mouse IgG antibody (Jackson ImmunoResearch 115-065-071) in 50% glycerol and 50% water was diluted to 1/500 in PBS-T with 2% (w/v) HI-FBS and incubated in wells for two hours at room temperature. Plates were then washed with PBS-T and Horseradish Peroxidase (HRP)-conjugated Avidin D (Vector Laboratories A-2014) diluted 1/1000 in PBS-T with 2% HI-FBS was added to each well for one hour at room temperature. Plates were then washed with PBS-T and then with 1X PBS. AEC substrate (BD Biosciences 551951) was added to each well, and plates placed in the dark for approximately five minutes. Plates were then rinsed in cold tap water, and the backing removed and rinsed. Plates were stored protected from light to dry and read within two working days.

#### Analysis

##### Neutralization assay

Absorbance was read at 405 nm. The average absorbance of background (cells only wells) was subtracted from each psV or psV+sample well absorbance prior to averaging signal in psV only wells and determining percent neutralization for each sample:

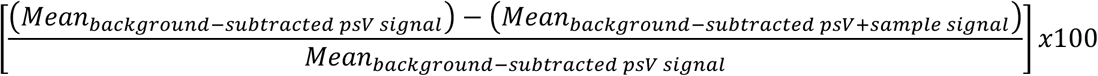

If the maximum average percent neutralization was lower than 50%, the sample was considered non-neutralizing and IC50 reported as half the reciprocal of the starting dilution (i.e., 50). Else, IC50 values were determined by fitting percent neutralization data curves using four parameter nonlinear regression (Prism v10).

##### ELISPOT

Plates were scanned using Autocenter on a CTL ImmunoSpot® S6 Universal Analyzer. Scans were counted using ImmunoSpot 5.3.22 software, Basic Count. Spot forming units (SFU) per well were normalized to the number of bone marrow cells added per well at each cellular dilution. SFU/well were averaged across all cellular dilutions of a given sample-antigen pair and then background corrected (i.e., the average SFU/well of corresponding cellular dilutions in PBS wells were subtracted). The frequency of HPV16- or HPV18-specific plasma cells was reported as a percentage of IgG-secreting plasma cells.

