## Supplemental Figures for "The magnitude and durability of neutralizing antibody responses to human papillomavirus vaccine do not depend on DNA sensing pathways"

#### Supplementary Figure 1

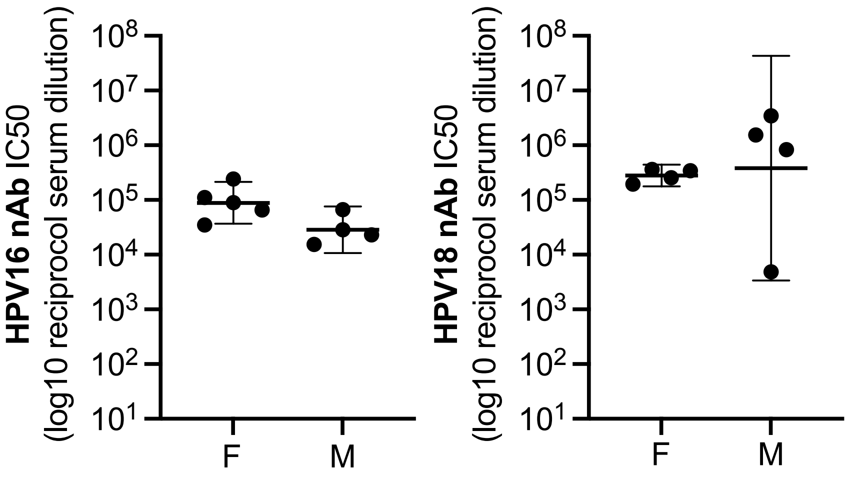


**Supplementary Figure 1 (related to Figure 1) Peak serum HPV16 and HPV18 nAb titers approximately one month after the third and final dose at Week 16 in mice that received 60µL 9vHPV vaccine dose volumes, stratified based on female (F) or male (M) sex as a biological variable.** Geometric mean with 95% confidence intervals shown, where each point represents the result from an individual mouse; Mann-Whitney U test between F and M.

#### Supplementary Figure 2

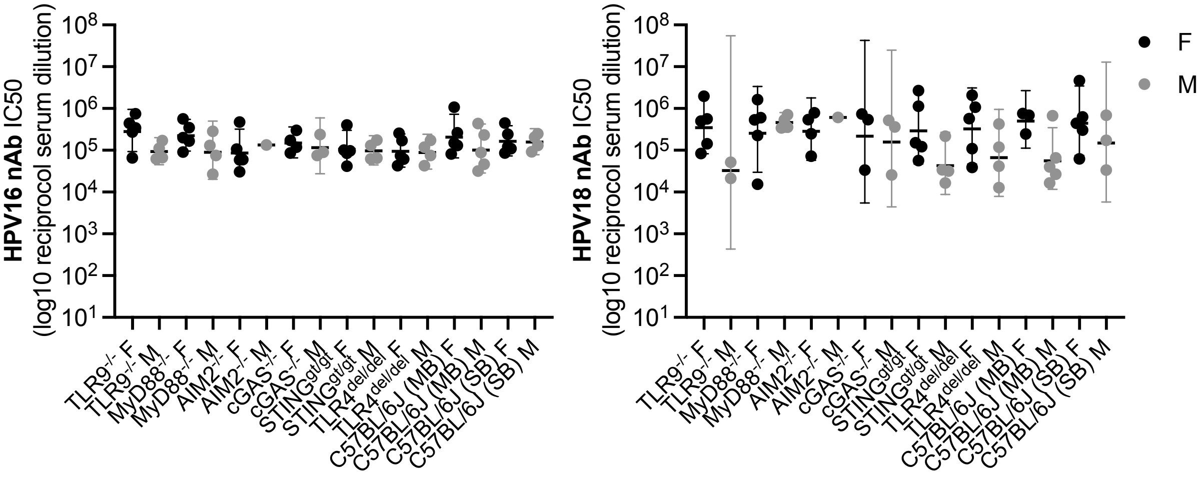


**Supplementary Figure 2 (related to Figure 2) Peak serum HPV16 and HPV18 nAb titers approximately one month after the third and final dose at Week 16 in mice that received 9vHPV, stratified based on sex as a biological variable.** Geometric mean with 95% confidence intervals shown, where each point represents the result from an individual mouse; Kruskal-Wallis with Dunn's post-test comparing F or M of similarly housed mice strains to F or M of control mice, respectively; Mann-Whitney U test between F and M of a given strain.

#### Supplementary Figure 3

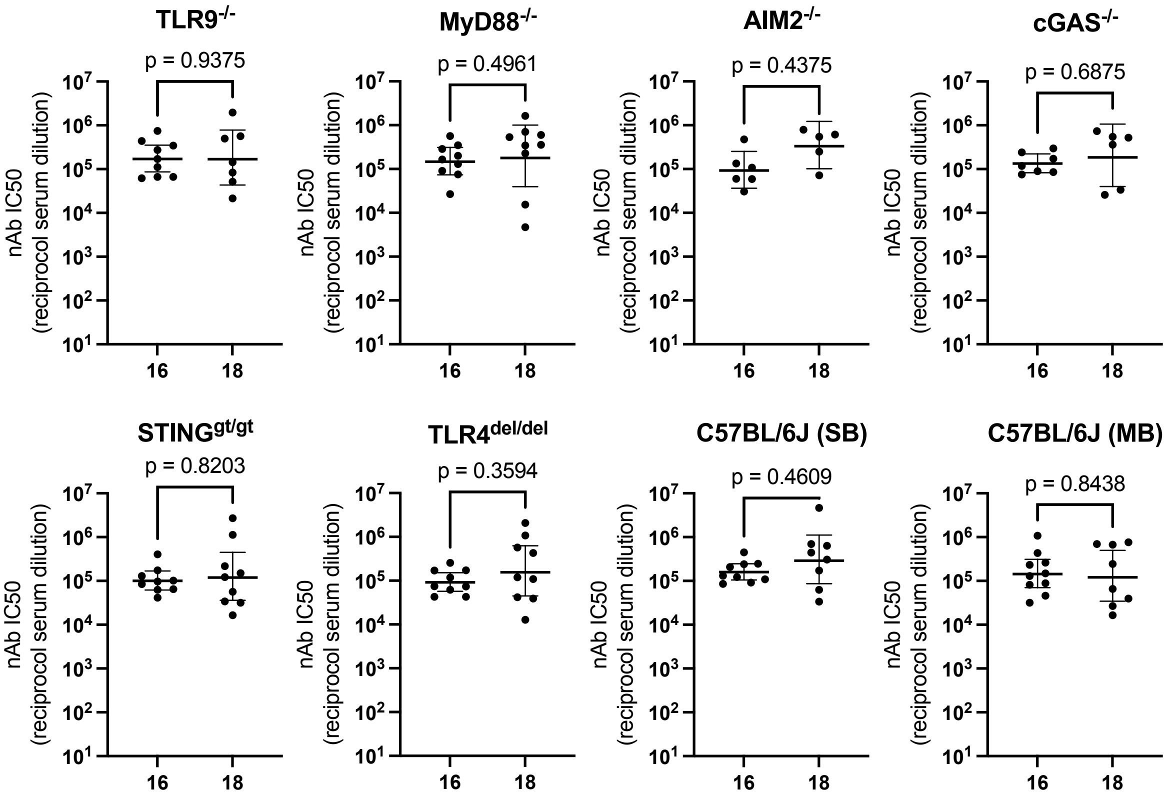


**Supplementary Figure 3 (related to Figure 2) No significant difference between peak serum HPV16 or HPV18 nAb titers one month after the third and final dose in different mice strains that received 9vHPV** Geometric mean with 95% confidence intervals shown, where each point represents the result from an individual mouse; Wilcoxon signed-rank test comparing HPV16 or HPV18 nAb GMT for a given strain.

#### Supplementary Figure 4

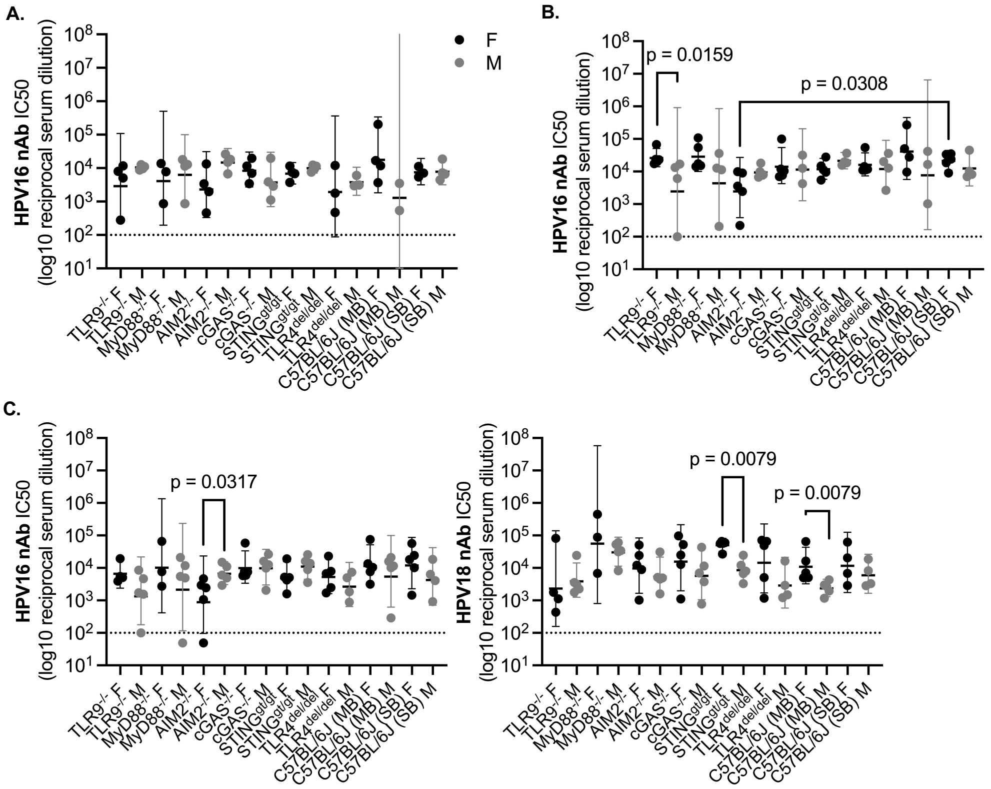

**Supplementary Figure 4 (related to Figure 3) Long-term serum HPV16 or HPV18 nAb titers in mice that received 9vHPV, stratified based on sex as a biological variable. A-B.** Serum HPV16 nAb titers approximately six months after the first 9vHPV vaccine dose at Week 26 (**A**) or approximately nine months after the first 9vHPV vaccine dose at Week 40 (**B**). **C.** Serum HPV16 and HPV18 nAb titers approximately one year after the first 9vHPV vaccine dose at Week 53-56. Geometric mean with 95% confidence intervals shown, where each point represents the result from an individual mouse; Kruskal-Wallis with Dunn's post-test comparing F or M of similarly housed mice strains to F or M of wildtype mice, respectively; Mann-Whitney U test between F and M of a given strain.

### Supplementary Figure 5

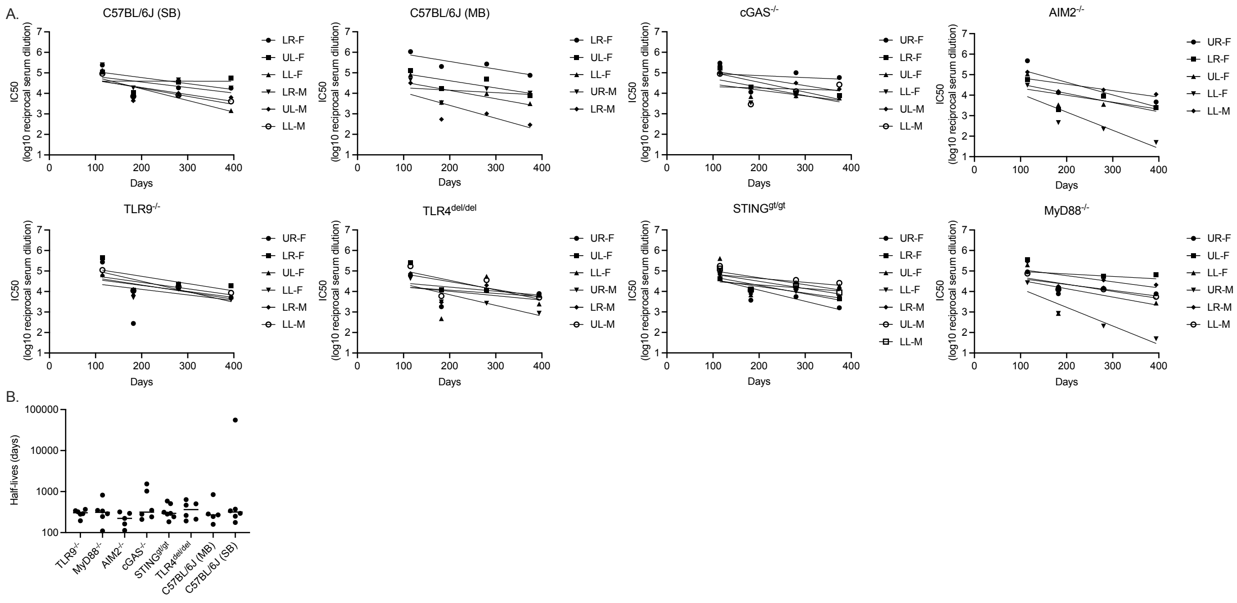


**Supplementary Figure 5 (related to Figure 3) Decay of HPV16 nAb titers over time in mice that received 9vHPV. A.** Serum HPV16 nAb titers in individual mice approximately four, six, nine, and twelve/thirteen months after the first 9vHPV vaccine dose. Curves fit with linear regression. Individual mice tags shown (e.g., LR-F = lower right ear tag, female mouse). C57BL/6J (SB), AIM2$^{-/-}$, TLR9$^{-/-}$, TLR4$^{del/del}$, and MyD88$^{-/-}$ mice euthanized at Week 56 (~13 months), whereas C57BL/6J (MB), cGAS$^{-/-}$, and STING$^{gt/gt}$ mice euthanized at Week 53 (~12 months). **B.** Serum HPV16 nAb titer half-lives per mouse strain as determined by linear regression in A. Median of each group shown, where each point represents the result from an individual mouse; Kruskal-Wallis with Dunn's post-test comparing similarly housed mice strains to wildtype mice.
